# Study on Vibrational Behavior of Cytoskeletons Modeled by Cylindrical Tensegrity Structure

**DOI:** 10.1101/2023.06.23.546353

**Authors:** Eiji Nouchi, Tomoteru Oka, Noriyuki Kataoka, Yoshihisa Kawano, Buntara Sthenly Gan

## Abstract

The dynamic mechanism of a cellular cytoskeleton is essential for the role of the cell, and its accurate characterization has been a long-standing problem for cell scientists. A cytoskeleton’s vibrations are highly influenced by interactions of filamentous proteins mediated by axial vibration of the stiff microtubules (compressive member) and lateral vibration of F-actin (tensile member). Among various structures in a cell, the cytoplasmic contractile bundles, lamellipodia, and filipodia cells can be modeled by a symmetrical cylinder-shaped self-equilibrium tensegrity structure with different radii at the top and bottom of the cylinder. The truncated conelike cylinder model is made to be small in height compared to both radii. This study investigates the tensegrity self-vibrational behavior of the cellular cytoskeleton to calculate its natural frequencies, composed of the individual vibration of microtubules and F-actins from measured data. The spectral element method is adopted based on the Wittrick–Williams procedure to solve the vibrational behaviors of the cellular cytoskeleton. Various n-polygon cylindrical truncated cone-shaped tensegrity structures to mimic the cellular cytoskeletons are presented to demonstrate the robustness of the present study.

## Introduction

Cells that makeup living organisms are exposed to various mechanical stimuli. In multicellular organisms, a network of cells forms a tissue. A system of tissues forms an organ (Deamer, 2007; Feher, 2017; Ingber, 1993a). Mechanical stimulation is involved in the background of cell morphology, motility, proliferation, and segregation and plays an important role in cell development and homeostasis. Inside a cell, there are organelles. Among the organelles, a cytoskeleton is a membrane that encloses the other organelles. The cellular cytoskeleton determines the cell’s shape and triggers cell movement and form changes (Albrecht-Buehler, 1987; Chicurel et al., 1998; Garder et al., 2008; Ingber et al., 1994; Ingber et al., 1995; Janmey 1998). When receiving an external force, stress is generated in the intracellular cytoskeleton and cell adhesive apparatus. It has become clear that proteins constituting the cytoskeleton and cell adhesive apparatus, such as actin filaments (actin filaments), perceive mechanical stimuli. The actin scaffold undergoes mechanical stimulation in the cytoskeleton and is rapidly remodeled (Stamenović, 2002). The role of this structure in mechanical response and its molecular mechanism has attracted attention (Feher, 2017; Pegoraro et al., 2017; Haspinger et al., 2021). Intermediate filaments have also been suggested to be involved in converting mechanical stimuli into chemical signals. However, although intermediate filaments are thought to play an important role in converting mechanical stimuli into chemical signals, the molecular mechanism remains unclear (Albrecht-Buehler, 1987; Dong et al., 1991; Fung and Liu, 1993; Janmey et al., 1991; Gittes et al., 1993; Schmid-Schönbein et al., 1995; Sieck, 2000).

In recent years, there has been an idea that the structure of a cytoskeleton is in the form of a tensegrity structure (Ingber et al., 1981; Ingber and Jamieson, 1982; Ingber, 1993b; Ingber, 2003a, 2003b). Although tensegrity was originally used in the self-equilibrium concept in the field of structural mechanics, its structure is believed to be applied to the shape of a cell. The cytoskeleton comprises actin filaments, intermediate filaments, and microtubules, forming a three-dimensional filamentous structure inside the cell (Fey et al., 1984; Heuser and Kirschner, 1980). Microtubules in the cytoskeleton are rigid, and actin and intermediate filaments are elastic, indicating a specific composition of a tensegrity structure.

A tensegrity structure is a statically indeterminate structure composed of continuous cables, tensile members, and discontinuous struts, compressive members, composite structures (Fuller, 1961, 1965; Snelson, 1996). Pins join these members, and the structure is in a self-equilibrium state without being supported. Many research results on tensegrity have been published in mechanics and architecture (Furuya, 1992; Motro et al., 1986). Various methods have been proposed in architecture for tensegrity structures’ morphogenesis and their vibration characteristics (Gan, 2019, 2022; Gan et al., 2015; Motro et al., 1986; Moussa et al., 2001; Murakami, 2001; Oppenheim and Williams, 2001; Sultan et al., 2002).

In this study, as an initial attempt, we imitate the vibrational behaviors of a cytoskeleton by using a twisted and truncated *n*-plex cylindrical tensegrity structure. Fig. 1 illustrates the tensegrity model used to mimic the cytoskeleton structure. Sensitivity analyses are conducted to investigate the parameters that rule the vibrational behaviors of the cytoskeleton. In a real application, the parameters can be calibrated by using a result obtained from measurements of the real cytoskeleton.

**Fig. 1.**
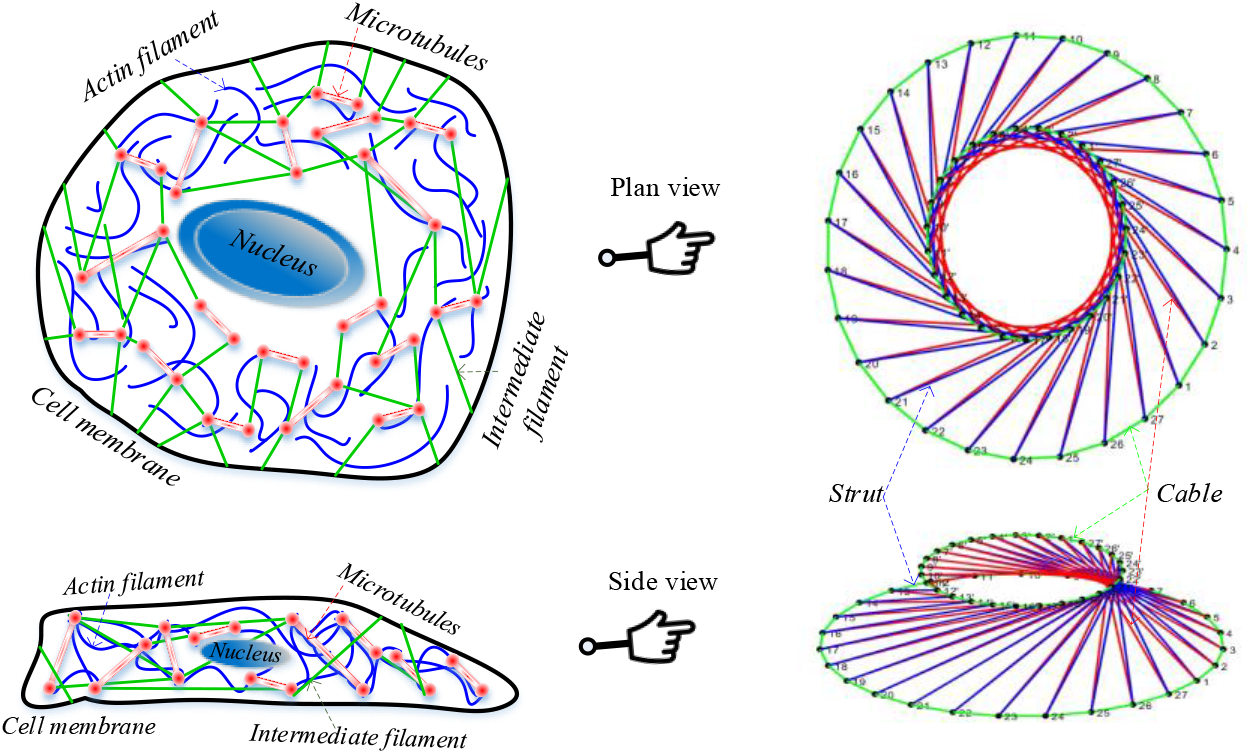
A twisted truncated polygon cylindrical tensegrity model to mimic a cytoskeleton structure. An illustration of a cellular cytoskeleton which is a twisted truncated polygon cylindrical tensegrity model.

### Truncated-cone tensegrity form

For illustration purposes, a quadruplex truncated cone (T-cone) can be constructed from 8 nodes and 16 members consisting of struts and cables, as shown in Fig. 2. Four nodes on the bottom and top of the cone form squares at the top and bottom.

**Fig. 2.**
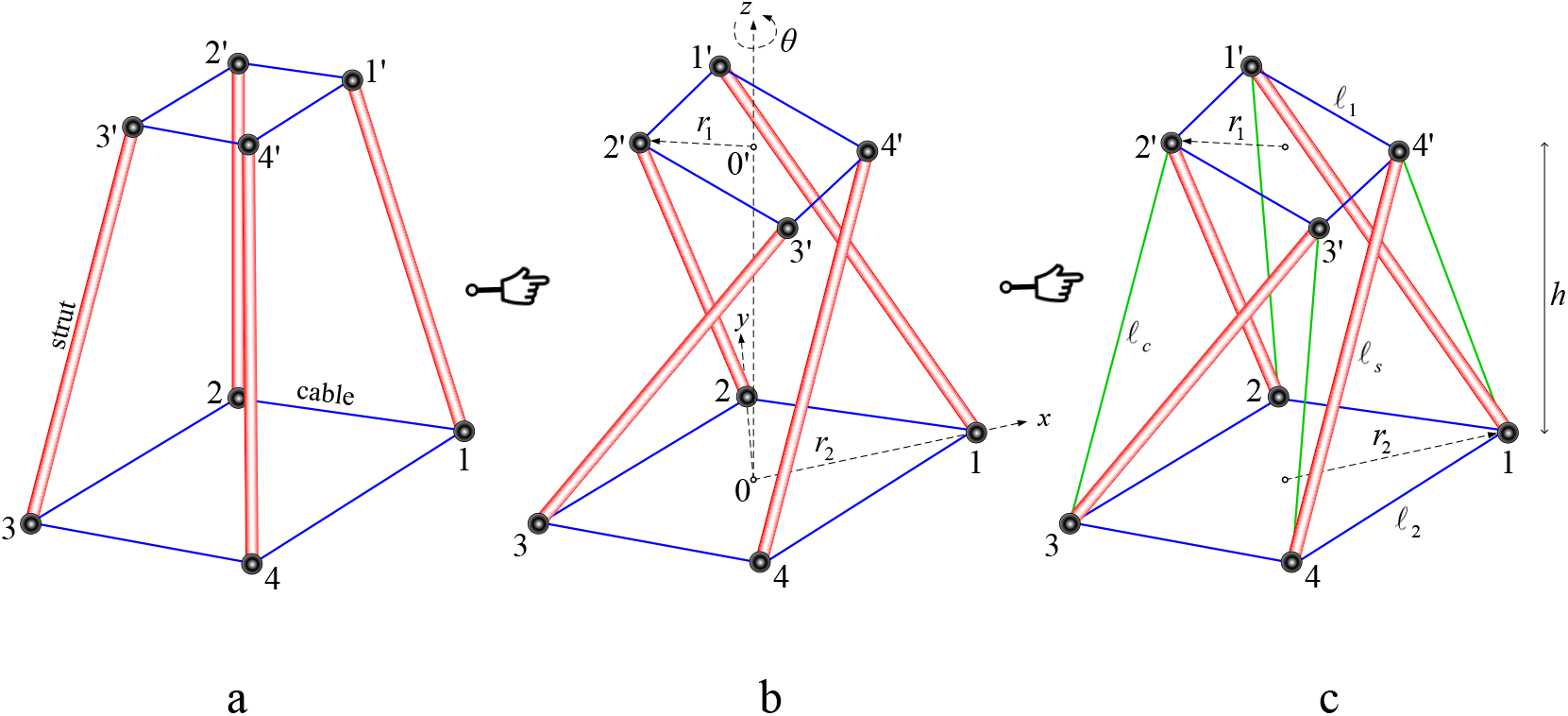
A twisted quadruplex T-cone tensegrity formation. Fig. 2a shows a quadruplex truncated conelike tensegrity model consisting of four struts (red) and eight cables (blue) of two different lengths (bottom and top). Fig. 2b shows the truncated conelike tensegrity being twisted by θ degrees at the center of the cone. Fig. 2c shows the additional four diagonal cables (green) to yield the self-equilibrium state of the tensegrity.

### Calculating the nodal coordinates

For constructing a T-cone-shaped tensegrity structure, a cylindrical coordinate system is the most convenient coordinate system for doing the analysis. Fig. 2 shows how the T-cone is twisted along the *z*-axis of both centers (made up of four nodes on a plane) 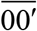 of the top and bottom squares. The center 0 of the bottom square lies at the origin of the coordinate system, while the center 0′ of the top square lies on the positive *z*-axis. The nodes which make up the square about the origin are marked by the labels 1, 2, 3, and 4. On the square at the top (height *h*) of the cone, the corresponding nodes are labeled 1′, 2′, 3′, and 4′. Thus, the nodes on the plane are equidistant (*r* is the radius of the circle) from the *z*-axis. The value of *θ* is the twisted angle of both squares.

Because nodes 1′, 2′, 3′, and 4′ also lie on the square rectangle, the angular components for nodes 2′, 3′, and 4′ are 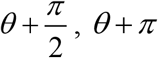, *θ* + *π*, and 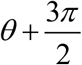, respectively. Table 1 summarizes the coordinates of the eight nodes of the quadruplex T-cone tensegrity structure.

**Table 1.** Polar coordinates of nodes in a quadruplex T-cone tensegrity structure.

| Node | $z$ | Radius | Angle |
| --- | --- | --- | --- |
| 1 | 0 | $r_1$ | 0 |
| 2 | 0 | $r_1$ | $\frac{\pi}{2}$ |
| 3 | 0 | $r_1$ | $\pi$ |
| 4 | 0 | $r_1$ | $\frac{3\pi}{2}$ |
| $1'$ | $h$ | $r_2$ | $\theta$ |
| $2'$ | $h$ | $r_2$ | $\theta + \frac{\pi}{2}$ |
| $3'$ | $h$ | $r_2$ | $\theta + \pi$ |
| $4'$ | $h$ | $r_2$ | $\theta + \frac{3\pi}{2}$ |

Let us put the compressive members of struts into the quadruplex T-cone tensegrity structure. The struts are placed at the line segments of 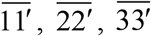, and 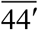. Next, the side bracing cables are specified to link up the two struts at the bottom and top of the cone. In Fig. 2, the side cables connect the line segments of 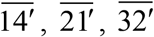, and 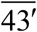.

We do an optimization algorithm to find the rotating angle *θ* so that the tensegrity structure is in the most 𝓁_*c*_ stable condition. The optimization is done by minimizing the lengths of bracing cables under the following constraining conditions:

- Constant radius *r*_1_ and *r*_2_ of the square rectangles of the cone *𝓁*_*s*_
- Constant length of the struts
- Symmetrical constraints of the struts 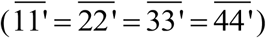
- Symmetrical constraints of the bracing cables 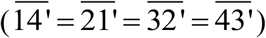

The general formulation for the length of the bracing cable can be given by

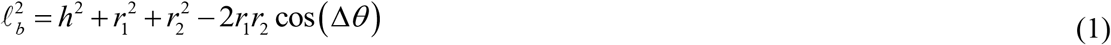

where Δ*θ* is the difference in angular coordinates between the two points.

The optimization problem can then be formulated as

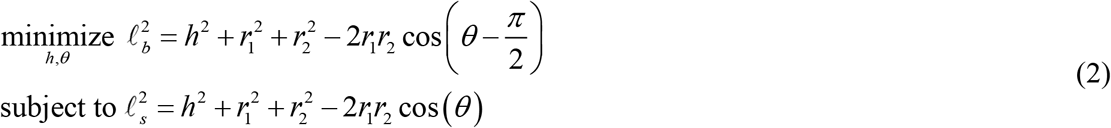

This constrained optimization problem can be transformed into an unconstrained one by using a constant radius ratio of 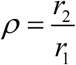, then by solving the constraint for the 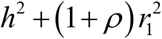 term from the second equation and substituting this term into the objective function. The unconstrained problem can now be defined as

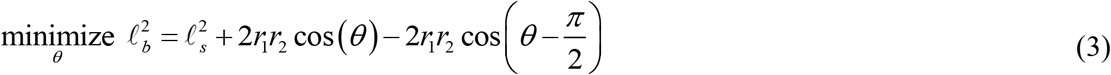

Taking the derivative with respect to *θ* and equating the result to zero yields:

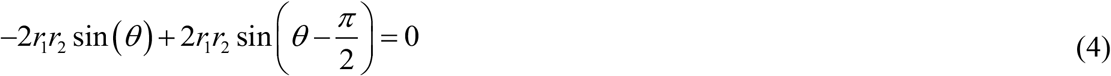

or

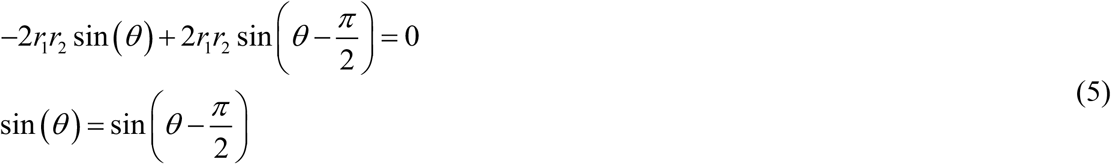

The sine of an angle has equality as follows:

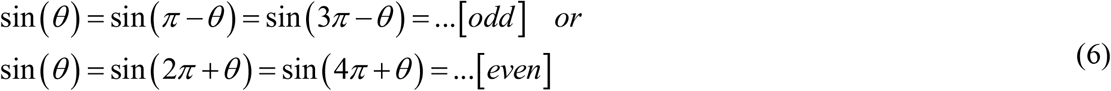

Only the odd multiplicity of *π* in (6) can give the solution to (5)

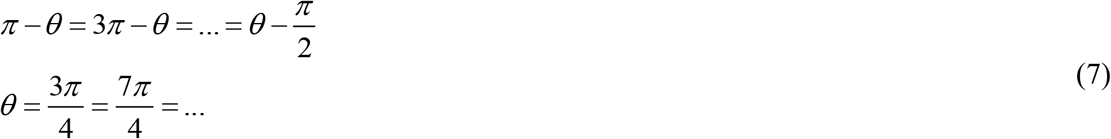

Substitution of the first term of 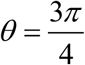 into (2) yields

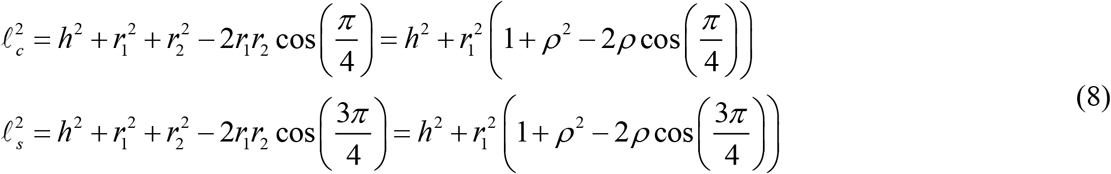

Assuming the length of the squared rectangular cable 𝓁_1_ = 1, introducing the radius of the constant radius ratio *ρ* = 2 and the length of the strut 𝓁_*s*_ = 10, then we can compute the radius *r*_1_, height *h*, and the length of the bracing cable 𝓁_*b*_ as

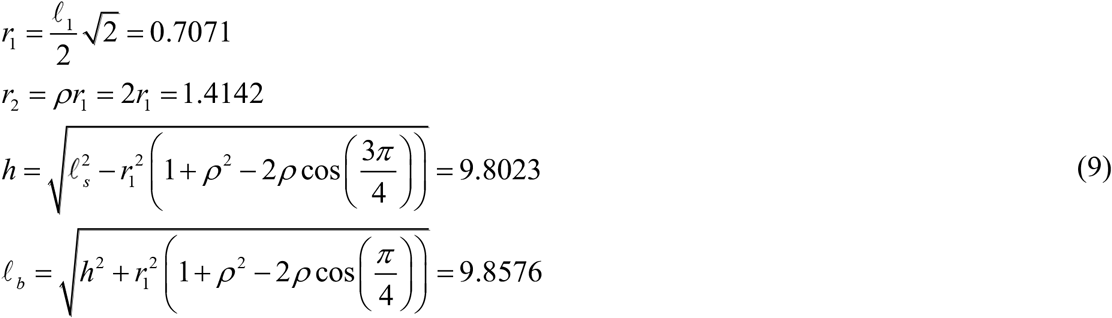

By substituting all the variables, the Cartesian coordinates of the nodes in the T-cone tensegrity structure can be summarized in Table 2.

**Table 2.** Cartesian coordinates of nodes in a quadruplex T-cone tensegrity structure.

| Node | $x$ | $y$ | $z$ |
| --- | --- | --- | --- |
| 1 | 1.4142 | 0.0000 | 0.0000 |
| 2 | 0.0000 | 1.4142 | 0.0000 |
| 3 | -1.4142 | 0.0000 | 0.0000 |
| 4 | 0.0000 | -1.4142 | 0.0000 |
| 1' | -0.5000 | 0.5000 | 9.8023 |
| 2' | -0.5000 | -0.5000 | 9.8023 |
| 3' | 0.5000 | -0.5000 | 9.8023 |
| 4' | 0.5000 | 0.5000 | 9.8023 |

### Calculating the tension coefficients

Because the T-cone tensegrity structure is fully symmetric, the equilibrated tension coefficients at node 1 also apply to those at all other nodes. In Fig. 3, it can be seen that node 1 is in equilibrium with nodes 2, 4, 1′ and 4′. The tension coefficients of the cables 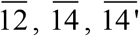, and 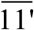 struts connected to node 1 are denoted by *q*_2_ = *q*_12_ = *q*_14_ (rectangular cables), *q*_*c*_ = *q*_14’_ (bracing cables), and *q*_*s*_ = *q*_11’_ (struts).

**Fig. 3.**
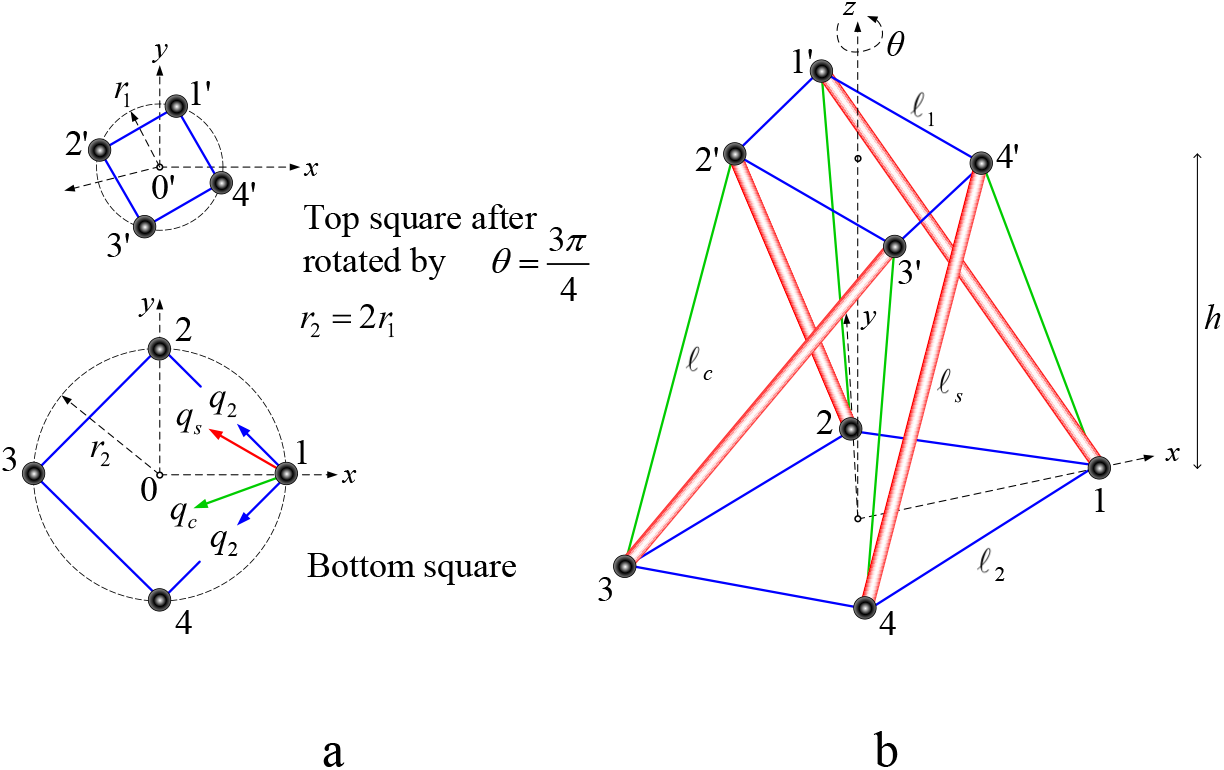
Equilibrium state of forces at node 1 in a quadruplex tensegrity structure. Fig 3a shows the plan views at the top and bottom squares of the tensegrity. The force equilibrium scheme is shown by vectors of forces from the strut and cables connected in node 1. Fig 3b shows the twisted and truncated quadruplex tensegrity structure.

Under the condition of no external loading applied, the equilibrium equations at node 1 in the *x*-, *y*- and *z*-directions can be obtained by substituting the nodal coordinates in Table 2 into the equations of force equilibrium at node 1 (Gan, 2019) becomes,

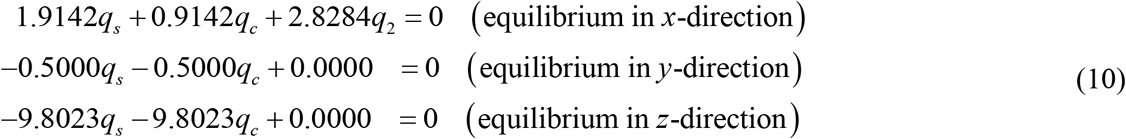

It can be verified that the equilibrium in *y*- and *z*-directions yields *q*_*c*_ = *−q*_*s*_. Assuming that the tension coefficients of the squared rectangular cables on the *x*–*y* plane are unitary, i.e. *q*_2_ = 1, the tension coefficients of the struts and the bracings are scalable to a non-zero constant without affecting the self-equilibrium state of the tensegrity structure. If we hold the compression strut force density as the variable *q*_*s*_ = *q*, the tension bracing cable density force becomes *q*_*c*_ = *−q*. Hence, the equilibrium in the *x*-direction of (10) can be written by

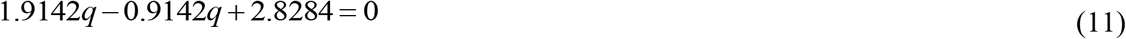

The tension coefficient *q* of the struts can be obtained as

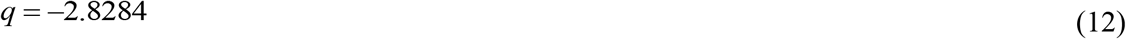

Similarly, the tension coefficient *q*_*c*_ in the bracing cables becomes

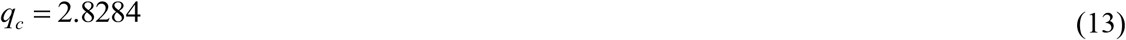

The tension coefficients of the squared rectangular cables on the top of the T-cone tensegrity structure can now be computed using the equilibrium in the *x*-direction at any node of choice, suppose at the node 1′ ; we can obtain the following equation.

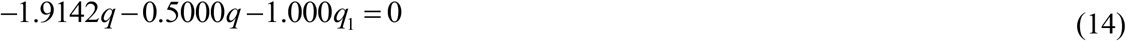

By substituting the tension coefficient *q* into the above equation, we can get

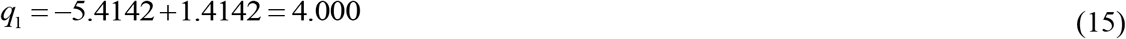

It can be observed that because the lengths of the top squared rectangular cables are half those of the bottom ones, the tension coefficient becomes four times larger than the bottom.

The determination of force coefficients of the members of the quadruple tensegrity structure above can be easily adapted to generate an *n*-polygon tensegrity structure. The force coefficients of the members are necessary to analyze the vibration of a tensegrity structure.

### Formulation of tensegrity structure in vibration

Fig. 4 shows a strut and a cable, two basic types of tensegrity members. The compressive member strut and tensile member cable represent the microtubule and actin filament, respectively. The vibration of the strut is modeled as an axially vibrating microtubule of a cytoskeleton. At the same time, the cable’s vibration is modeled to be a laterally vibrating actin filament of a cytoskeleton. The microtubule is presumed to have an axial rigidity represented by prestress force, *P*, and Young’s modulus of elasticity, *E*. In contrast, the actin filament is presumed to have only an axial rigidity represented by prestress force, *P*.

**Fig. 4.**
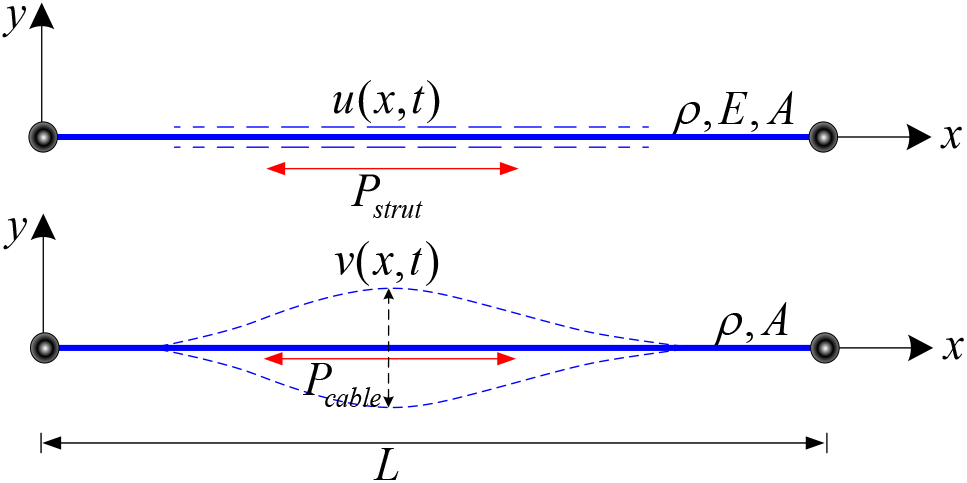
Vibrational models of strut and cable members. The vibration of a strut member is assumed to be in the axial direction of the member axis. Because the cable cannot vibrate in the axial direction of the axis, the vibration of a cable member is considered to be in the lateral direction of the axis.

The equations of vibrational motion (Gan, 2022) are given as follows:

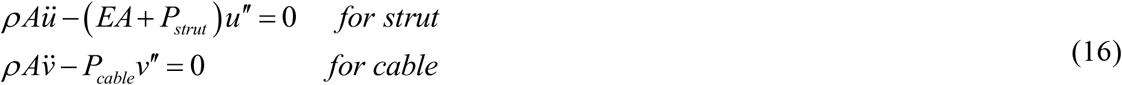

where *u*(*x,t*) and *v*(*x,t*) are the axial and lateral displacements which are a function of position and time, respectively. The dot and prime superscripts denote the derivatives with respect to time *t* and spatial coordinate *x*, respectively.

The internal forces and boundary conditions are given as

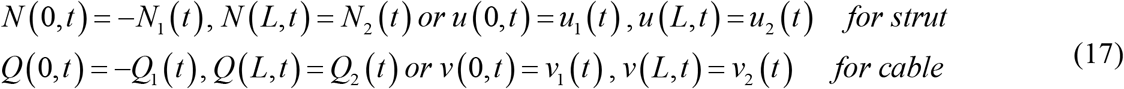

where *N*(*x,t*), *Q*(*x,t*) are the axial and shear forces defined by

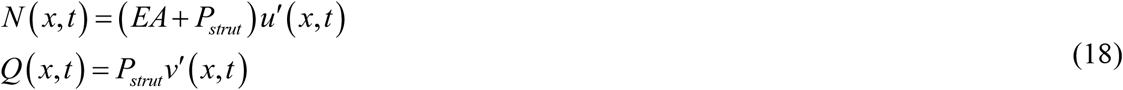

### Spectral element formulation

The analysis of the vibrational behaviors of the tensegrity was performed by using the Spectral Element Method (SEM, Beskos and Narayanan, 1983; Lee, 2009), which is a combination of the Finite Element Method (FEM), the Dynamic Stiffness Method (DSM, Banarjee, 1997; Leung, 1993), and the Spectral Analysis Method (SAM, Doyle, 1989). The vibration of a tensegrity structure varies with the individual vibrational frequency and the wavelength of the members.

The FEM is a famous computational method used in many fields of engineering and science. However, it is difficult for the conventional FEM to analyze a structure such as a tensegrity structure with no supporting boundaries, no external force, and members in a self-equilibrium state. The DSM uses an exact dynamic stiffness matrix and implicitly considers the mass in the stiffness equation. The SAM uses the Fast Fourier Transform and has the property that the error converges exponentially if the solution is a smooth function (“exponential convergence”), converging much faster than the FEM. In spectral analysis, the dominant differential equation can be solved by infinitely adding waves with different frequencies and the time history of the solution that can be obtained from the Inverse Fourier Transform in the frequency-domain spectral components.

Fig. 5 depicts the concept of the SEM as a combination of FEM, SAM, and SAM techniques in computing the vibration of tensegrity structures.

**Fig. 5.**
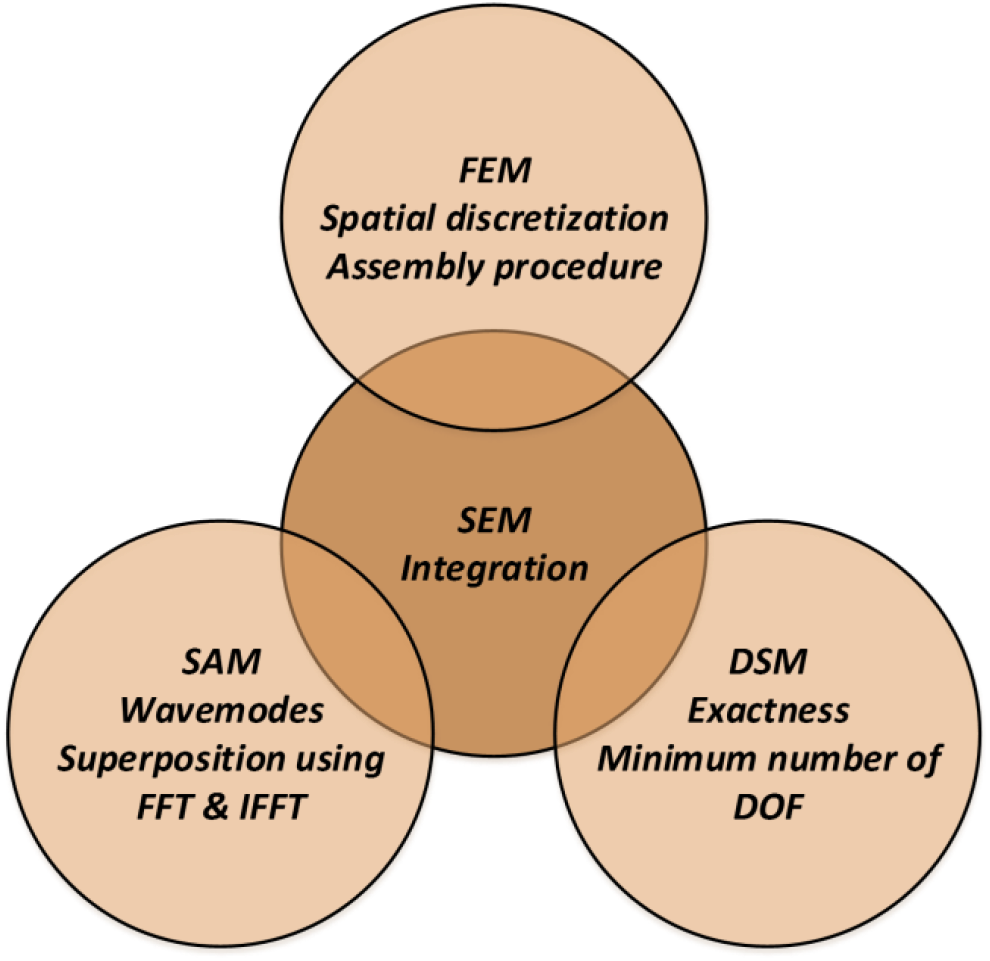
Outline of the Spectral Element Method (SEM). SEM is a new method combining the discretization feature of FEM, the exact solution solver of DSM, and the wave modes analysis using the Fourier Transform.

### Governing equations in the frequency domain

Assume the dynamic response of the tensegrity structure in generalized coordinate *w*(*x,t*), whereas in the spectral form given by

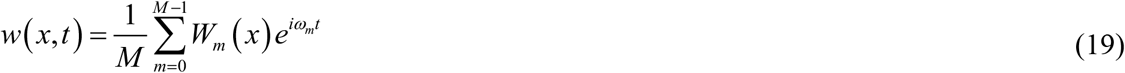

where 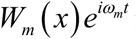 is the generalized spectral components of *u*(*x,t*) or *v*(*x, t*), *M* is the sampling number, and *ω*_*m*_ is the *m*^*th*^ natural frequency.

In spectral forms, the generalized boundary forces *F*_1_(*t*) and *F*_2_(*t*), and the generalized boundary displacements *w*_1_(*t*) and *w*_2_(*t*) are also assumed to be as follows:

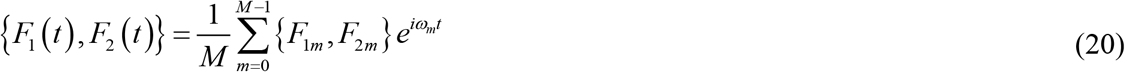

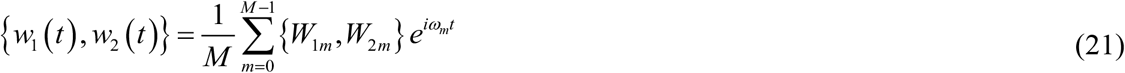

### Spectral nodal degrees of freedom (DOF) and forces

The spectral nodal generalized displacements *W*(*U, V*) and forces *F*(*N,Q*) are depicted in Fig. 6.

**Fig. 6.**
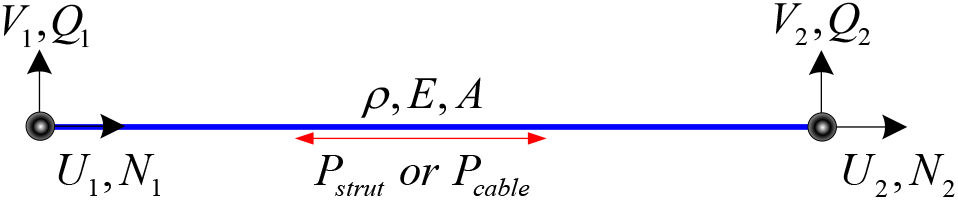
Spectral nodal generalized DOFs and forces. The strut and cable members are modeled using two nodes at both ends. At each node, there are two generalized degree of freedoms (horizontal and vertical) in the axis direction. At the node, these DOFs represent the direction of displacements (*U*_1_,*U*_2_, *V*_1_,*V*_2_) and their corresponding force reactions (*N*_1_, *N*_2_, *Q*_1_,*Q*_2_) at the nodes.

Detailed derivation of formulas can be found in [2]. Only relevant formulas are presented in the following.

The spectral nodal DOF vector is given by

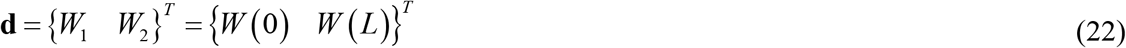

and the spectral nodal forces vector is given by

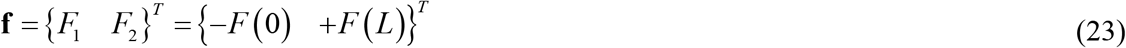

### Dynamic shape function

The solution of (19) can be given by

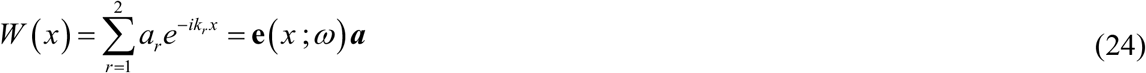

where the two wavenumbers can be determined by

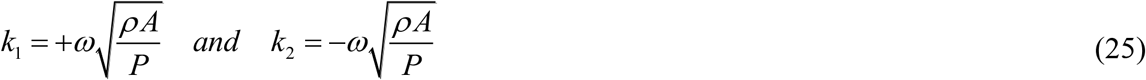

with 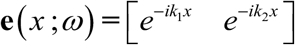 and ***a=*** {*a*_*1*_ *a*_*2*_}^*T*^.

The dynamic shape function can be obtained from

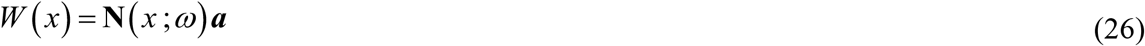

where **N**(*x* ;*ω*) is given by

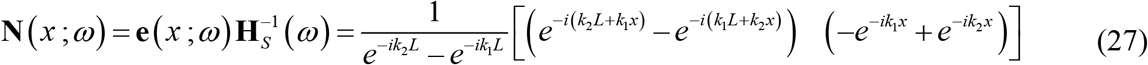

The spectral nodal DOF vector of (22) can be formulated by substitution of (26) into (27), which gives

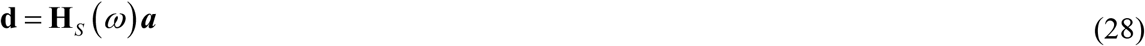

where

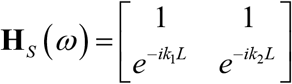

### Weak form of governing equation

The spectral element equation of tensegrity structures can be given as

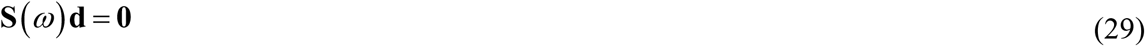

where

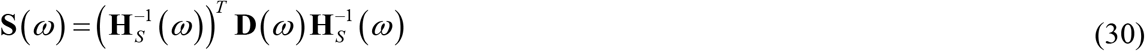

with

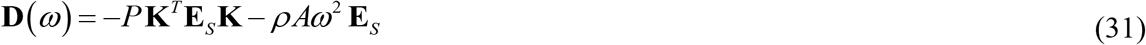

The derivation of the dynamic shape function (27) can be written as

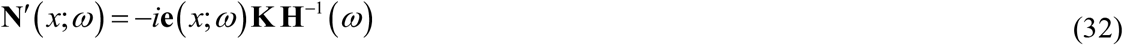

where

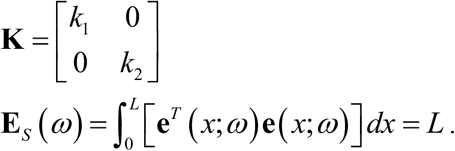

The Wittrick–Williams procedure is adopted to solve the natural vibrations of the cellular cytoskeleton based on the dynamic tensegrity formulations above (Wittrick and Williams, 1971).

### Vibrational characteristics of tensegrity structures

Analysis was performed using the SEM, a combination of the FEM, the DSM, and the SAM.

### Natural vibrations of components of tensegrity structures

The vibration of a tensegrity structure is determined by the combination of the vibration behaviors of its components (cables and struts). Physical data of each cytoskeleton component were obtained from the following literature (Demé et al., 2000; Feher, 2017; Haspinger et al., 2021; Igaev and Grubmüller, 2020; Perogaro et al., 2017; Gardel et al., 2008).

- Length: Actin = 3∼20 μm; microtubule = 10 μm
- Diameter: Actin = 7∼10 nm; microtubule = 25 nm
- Density: Actin = 1.38∼1.40 g/cm^3^; microtubule = 1.41 g/cm^3^
- Elastic modulus: Microtubule = ∼ 1 GPa

The tensions per unit length are obtained from the self-equilibrium equation (Gan, 2019). Table 3 shows the physical properties of 9-plex cytoskeleton components based on the literature.

**Table 3.** Physical properties of 9-plex cytoskeleton components.

| Component | Microtubule<br>(diagonal<br>strut) | Actin filament<br>(top and bottom<br>cables) | Actin filament<br>(diagonal cable) |
| --- | --- | --- | --- |
| Length (nm) | 1912.8 | 500.0 | 1711.1 |
| Diameter (nm) | 25 | 7 | 7 |
| Density ($\mu\text{g/nm}^3$) | 1.41e-15 | 1.38e-15 | 1.38e-15 |
| Elastic modulus (Pa) | 3.0e-5 | - | - |
| Tension per unit length (aN/nm) | -2.736e-6 | 4.0e-8 | 2.736e-6 |

Fig. 7 shows the model of a 9-plex cytoskeleton tensegrity structure and the results of natural frequencies that satisfy (29), as can be seen as drops in the logarithmic values of the determinant of dynamic stiffness. The first drop is called the structure’s first mode of natural frequency.

**Fig. 7.**
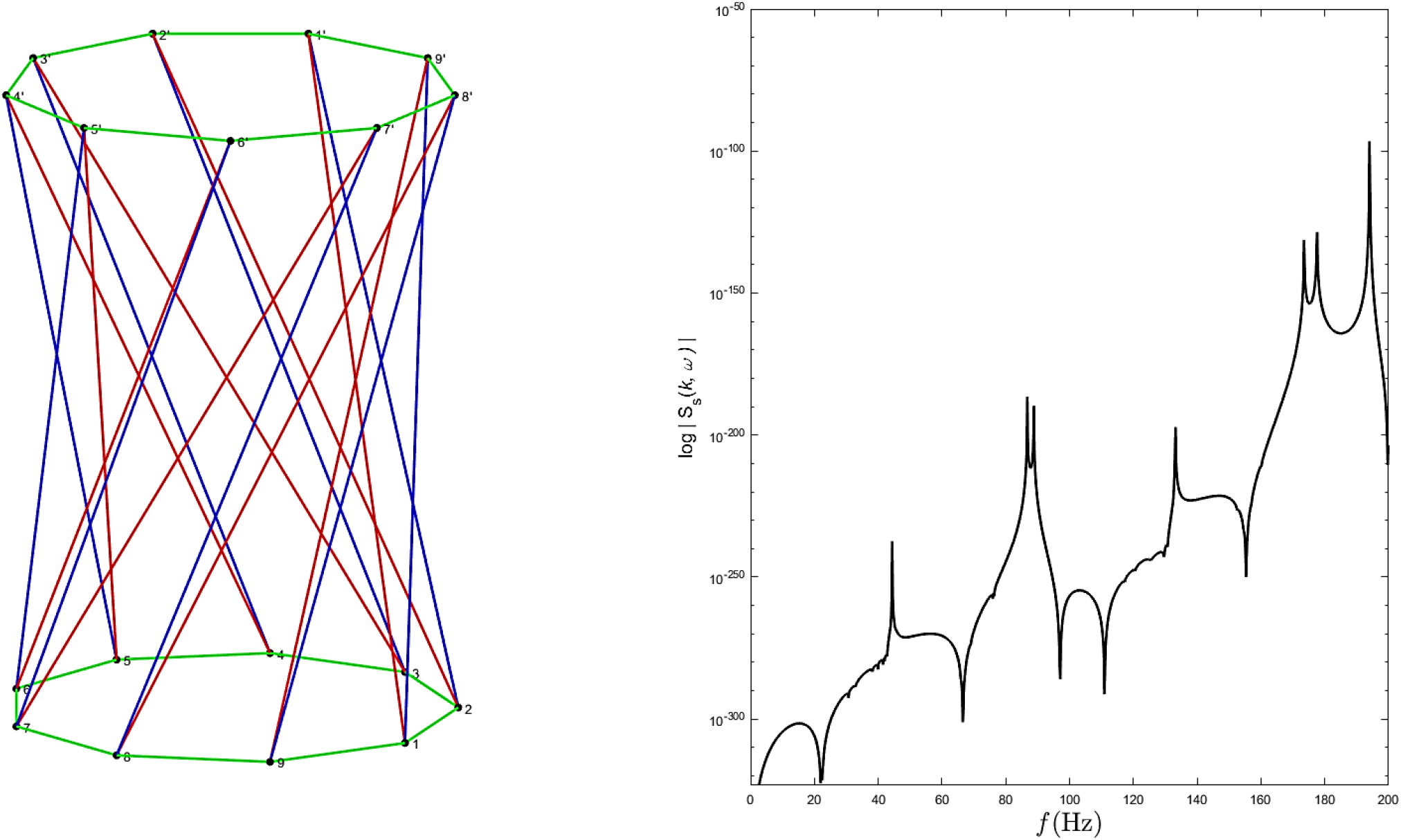
Vibration of a 9-plex cytoskeleton tensegrity structure (1^st^ mode: 21.963 Hz). The left side of Fig. 7 shows the 9-plex cytoskeleton tensegrity with similar radii at the bottom and top of the cylindrical. The Wittrick-Williams procedure is adopted to solve the natural frequency vibrations of the cellular cytoskeleton based on the dynamic tensegrity formulations.

### Height variation

The variation of heights of the cytoskeleton structure is investigated in this section. Table 4 shows the height variation configurations of the tensegrity structures with their physical properties, which are required to compute the natural frequencies of the structures.

**Table 4.** Physical properties of 9-plex tensegrities.

| Height (nm) |  | 1193.2<br>(0.2H<sub>0</sub>) | 894.94<br>(0.6H<sub>0</sub>) | 1193.2<br>(0.8H<sub>0</sub>) | 1491.6<br>(1.0H<sub>0</sub>) |
| --- | --- | --- | --- | --- | --- |
| Length (nm) | Microtubule | 1234.1 | 1495.0 | 1690.5 | 1912.8 |
|  | Actin top & bottom |  | 500.0 |  |  |
|  | Actin diagonal | 890.0 | 1226.4 | 1458.4 | 1711.1 |
| Diameter (nm) | Microtubule |  | 25 |  |  |
|  | Actin |  | 7 |  |  |
| Density<sup>3)</sup> (μg/nm<sup>3</sup>) | Microtubule |  | 1.41e−15 |  |  |
|  | Actin |  | 1.38e−15 |  |  |
| Elastic modulus (Pa) | Microtubule |  | 3.0e−5 |  |  |
| Tensile coefficient (aN/nm) | Actin top (as reference) |  | 4.0e−6 |  |  |

Figures 9–11 show the 9-plex cytoskeleton tensegrity structure model with height variations of 0.8H, 0.6H, and 0.2H, respectively. The first mode of natural frequencies of the various height cytoskeleton tensegrity structures that satisfy (29) can be seen in the first drops in the logarithmic values of the determinant of dynamic stiffness.

Fig. 8 shows natural frequency graphs of each 9-plex cytoskeleton tensegrity structure component.

**Fig. 8.**
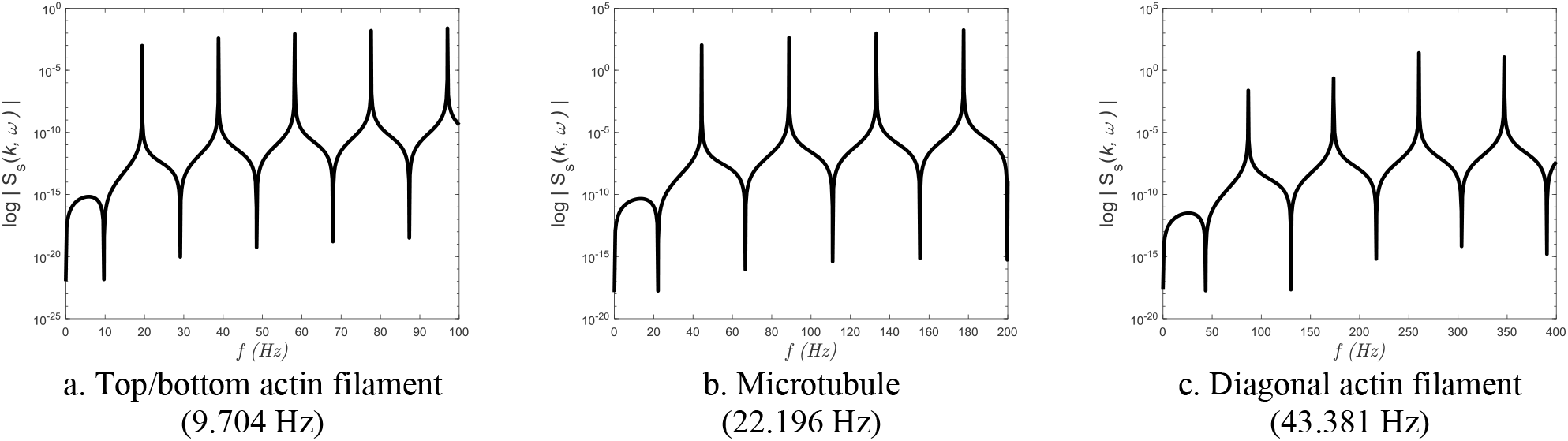
The first mode from five-mode vibration of struts and cables in a 9-plex tensegrity cell. Fig. 8a shows the first frequency of 9.704 Hz of a single actin filament modeled by the cables (green) in the tensegrity model. Fig. 8b shows the first frequency of 22.196 Hz of a single microtubule modeled by the struts (red) in the tensegrity model. Fig. 8c shows the first frequency of 43.381 Hz of a single actin filament modeled by the cables (blue) in the tensegrity model.

**Fig. 9.**
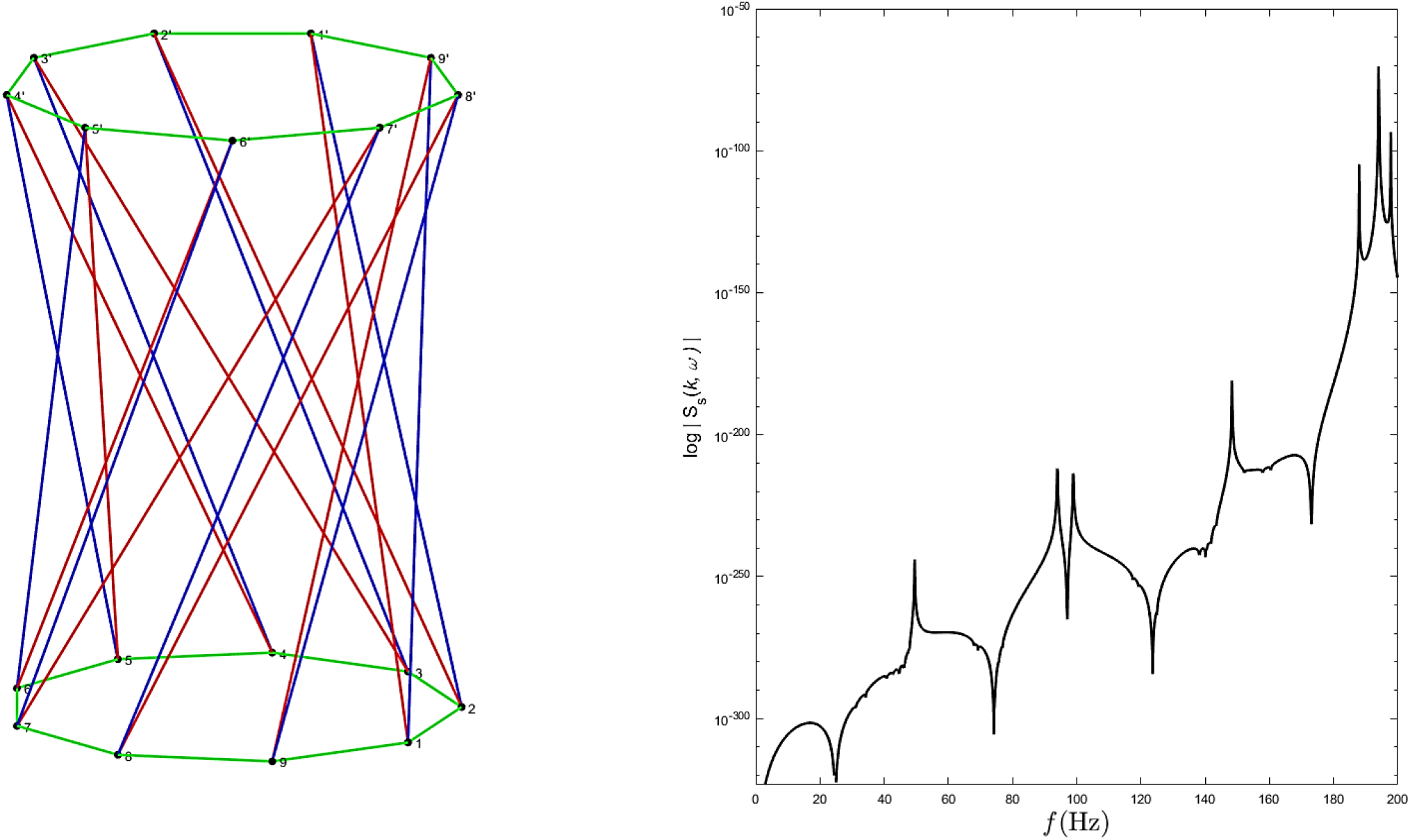
Vibration of a 9-plex tensegrity cell (0.8H = 25.000 Hz). The natural frequencies of the 0.8 times the initial height tensegrity when all the single members were composed show the first frequency of 25.000 Hz, which is higher (stiffer) than the single member of cables (actin) on the bottom and top of the tensegrity (see Fig. 8). However, the first natural frequency of the tensegrity is still lower (softer) than the individual diagonal member of cables (actin).

**Fig. 10.**
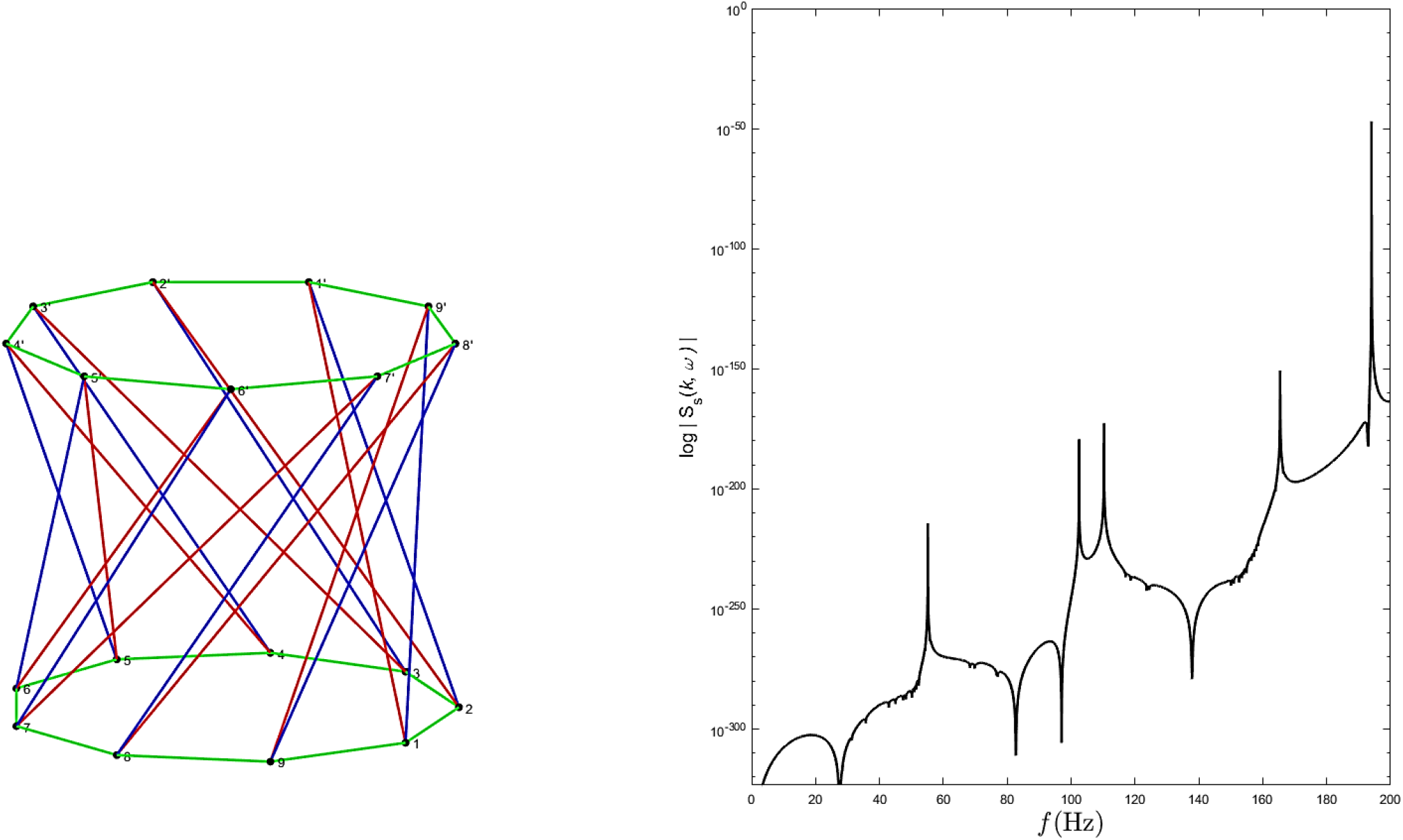
Vibration of a 9-plex tensegrity cell (0.6H = 27.399 Hz). The natural frequencies of the 0.6 times the initial height tensegrity when all the single members were composed show the first frequency of 27.399 Hz, which is higher (stiffer) than the single member of cables (actin) on the bottom and top of the tensegrity (see Fig. 8). However, the first natural frequency of the tensegrity is still lower (softer) than the individual diagonal member of cables (actin).

**Fig. 11.**
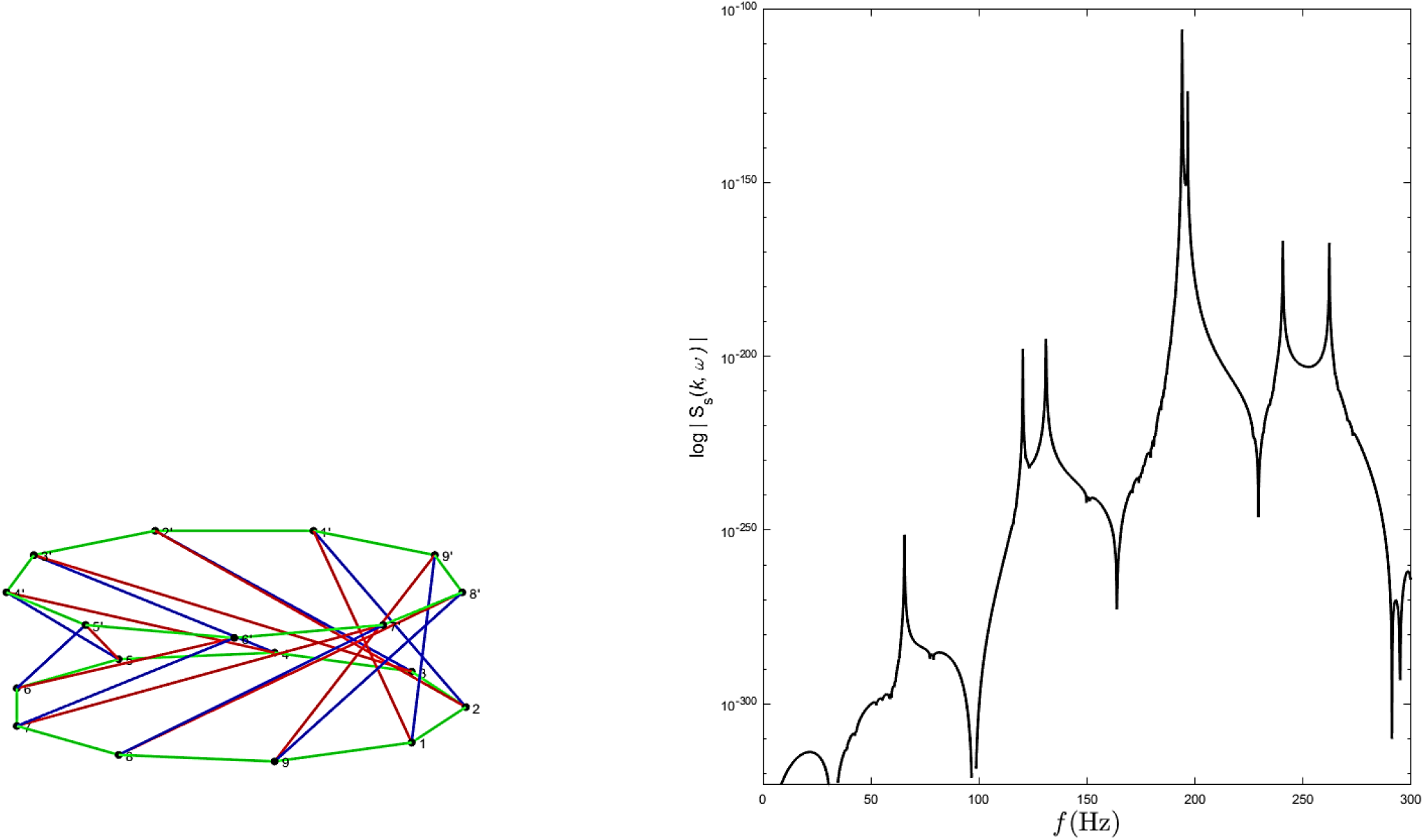
Vibration of a 9-plex tensegrity cell (0.2H = 30.240 Hz). The natural frequencies of the 0.2 times the initial height tensegrity when all the single members were composed show the first frequency of 30.240 Hz, which is higher (stiffer) than the single member of cables (actin) on the bottom and top of the tensegrity (see Fig. 8). However, the first natural frequency of the tensegrity is still lower (softer) than the individual diagonal member of cables (actin).

### n-Plex variation

This section investigates the variation of n-plex cytoskeleton twisted cylindrical tensegrity structures with a height of 1.4% of H_0_. Table 5 shows the physical properties of 9-, 12-, and 15-plex cytoskeleton tensegrity structures, respectively.

**Table 5.**
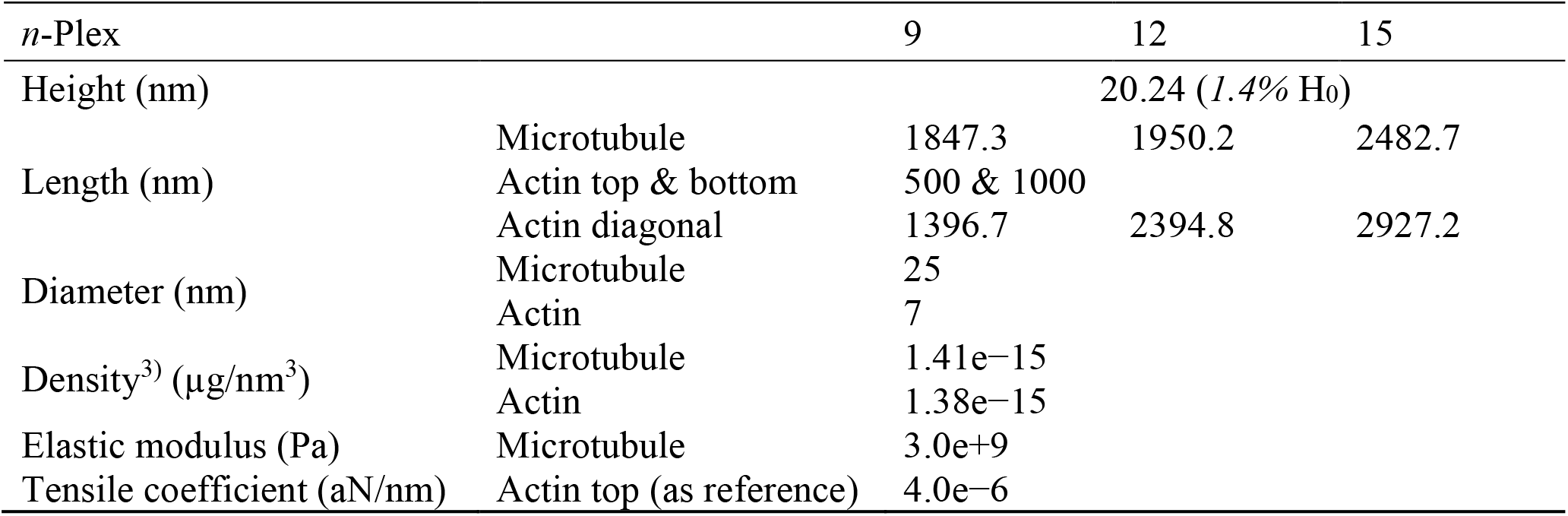
Physical properties of *n*-plex tensegrities.

| $n$-Plex | | 9 | 12 | 15 |
| --- | --- | --- | --- | --- |
| Height (nm) | | 20.24 (1.4% $H_0$) | | |
| Length (nm) | Microtubule | 1847.3 | 1950.2 | 2482.7 |
|  | Actin top & bottom | 500 & 1000 |  |  |
|  | Actin diagonal | 1396.7 | 2394.8 | 2927.2 |
| Diameter (nm) | Microtubule | 25 |  |  |
|  | Actin | 7 |  |  |
| Density<sup>3)</sup> ($\mu\text{g}/\text{nm}^3$) | Microtubule | 1.41e-15 | | |
|  | Actin | 1.38e-15 |  |  |
| Elastic modulus (Pa) | Microtubule | 3.0e+9 |  |  |
| Tensile coefficient (aN/nm) | Actin top (as reference) | 4.0e-6 |  |  |

Figures 12–14 show the model of an *n*-plex cytoskeleton tensegrity structure with a height of 0.2H. The first mode of natural frequencies of the *n*-plex cytoskeleton tensegrity structures that satisfy (29) can be seen in the first drops in the logarithmic values of the determinant of dynamic stiffness.

**Fig. 12.**
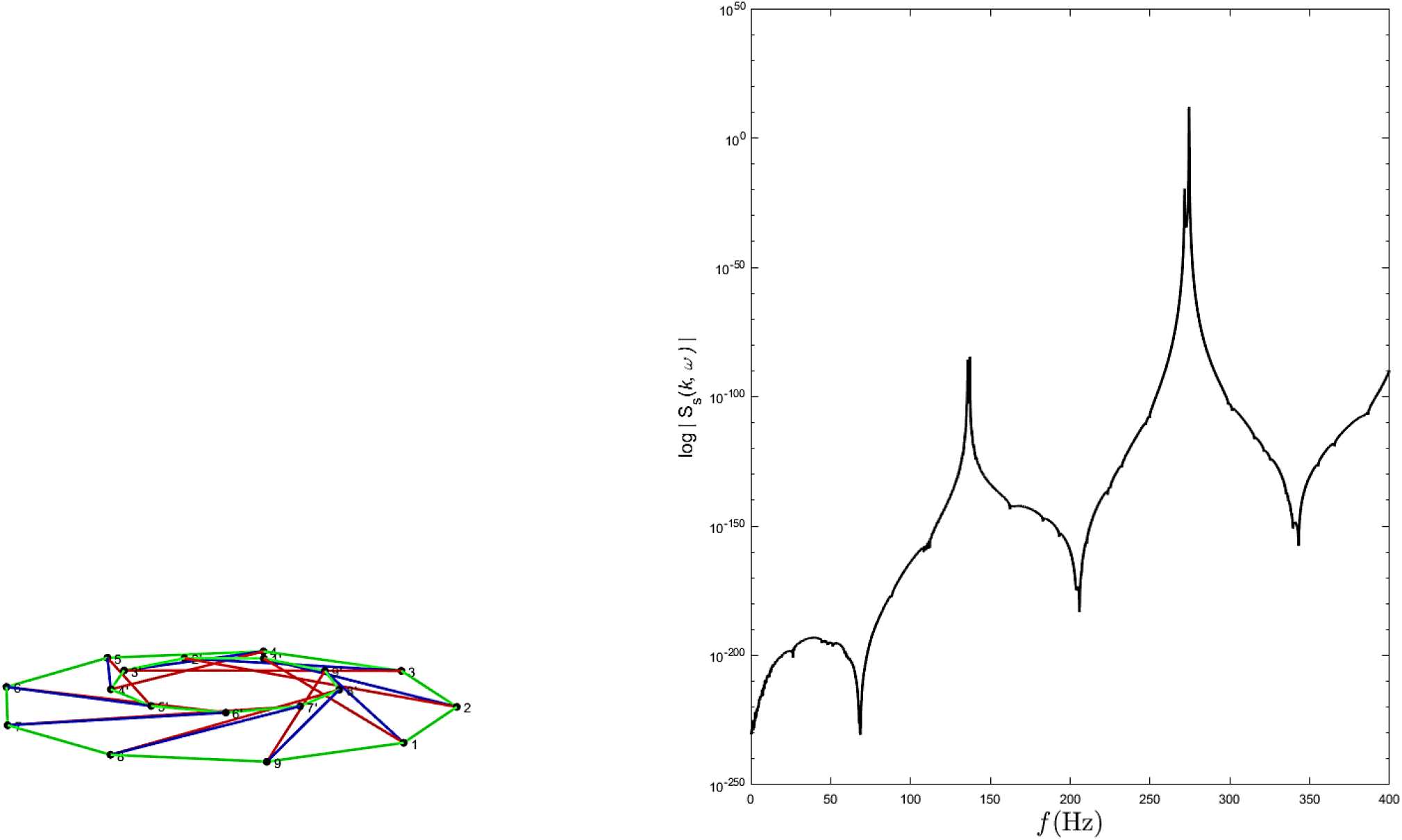
Vibration of a truncated 9-plex conic tensegrity cell (*f*_1_ = 68.61 Hz). The natural frequencies of the 1.4% initial height of the 9-plex tensegrity composed by individual members show the first frequency of 68.61 Hz, which is higher (stiffer) than all individual members.

**Fig. 13.**
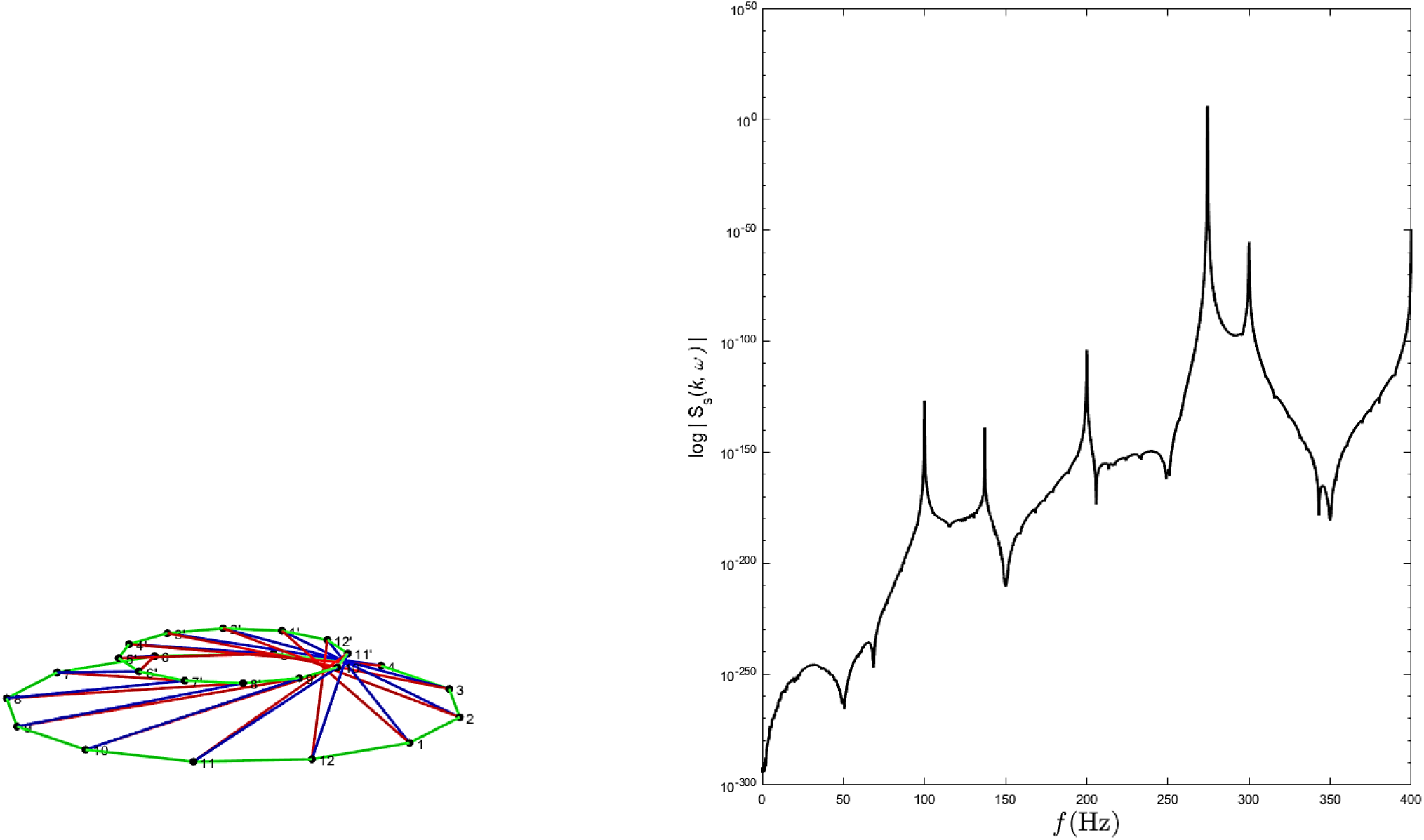
Vibration of a truncated 12-plex conic tensegrity cell (*f*_1_ = 50.61 Hz). The natural frequencies of the 1.4% initial height of the 12-plex tensegrity composed by individual members show the first frequency of 50.61 Hz, which is higher (stiffer) than all individual members.

**Fig. 14.**
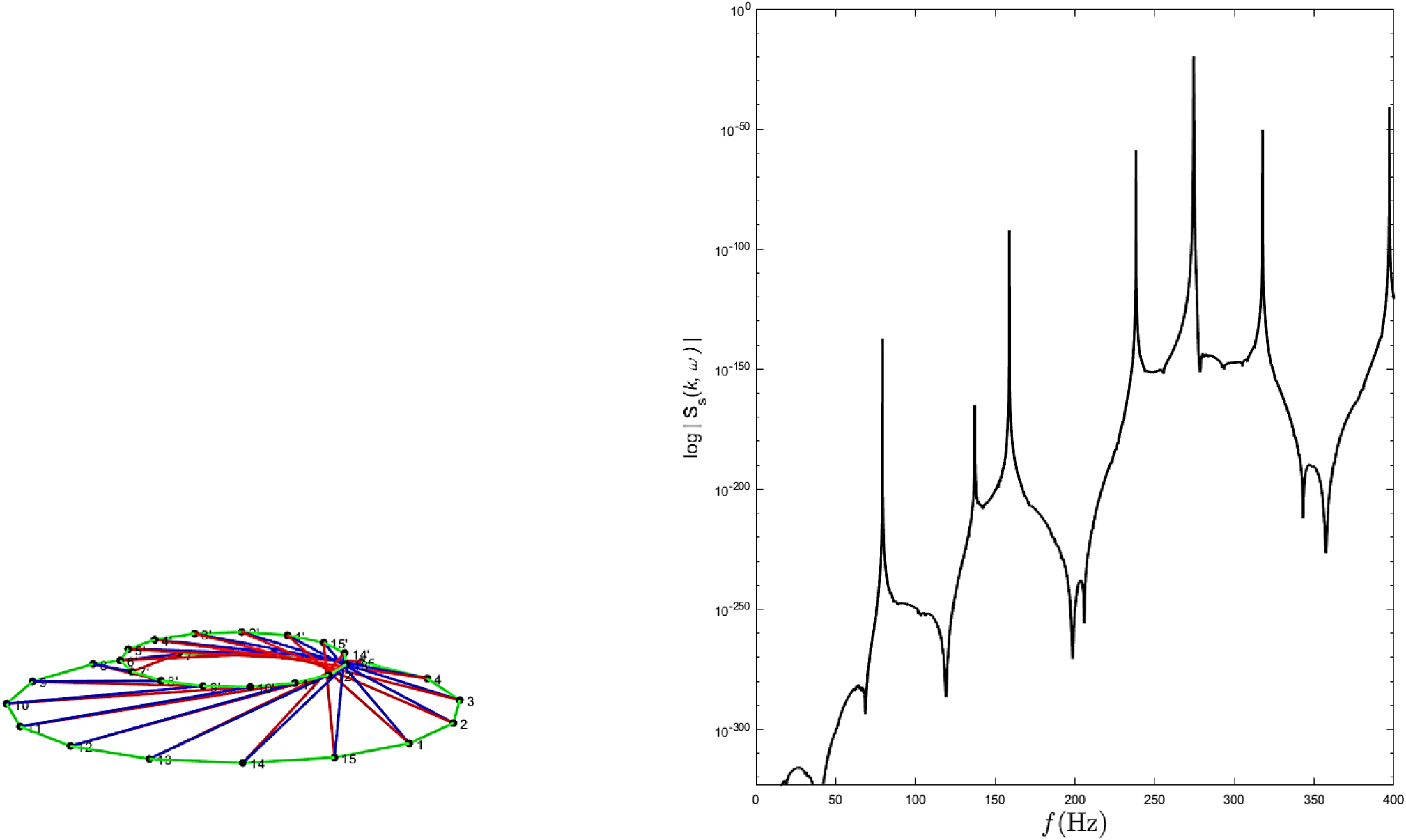
Vibration of a truncated 15-plex conic tensegrity cell (*f*_1_ = 35.49 Hz). The natural frequencies of the 1.4% initial height of the 15-plex tensegrity composed by individual members show the first frequency of 35.48 Hz, which is lower (softer) than the individual diagonal member of cables (actin) but higher (stiffer) than the other individual members in the tensegrity.

## Summary and Conclusion

In Figs. 7 and 8, we can observe that the cellular 9-plex tensegrity structure has a 1^st^ mode natural frequency which is close to the 1^st^ mode natural frequency of its strut (microtubule) because the actin filaments are assumed to vibrate in their lateral direction. In contrast, the microtubule is assumed to vibrate in its axial direction, controlling the vibrational behaviors of the tensegrity structure (Ghavanloo et al., 2010; Pokorný et al., 1997; Tounsi et al., 2010).

The height of a 9-plex tensegrity structure is varied by 0.8, 0.6, and 0.2 times the initial height H. Figs. 9–11 show that the 1^st^ mode of their natural frequency was increased by reducing the height.

Fixing the height of the 9-plex tensegrity at 1.4%H, the 1^st^ modes of the 12- and 15-plex tensegrity structures show an increasing natural frequency since they become stiffer than the 9-plex tensegrity.

Using tensegrity structures, we obtained physical properties to reproduce the fluctuations of cells. In the future, we will investigate the significance of fluctuations in the cellular structure composed of actin filaments and microtubules subjected to traction force by observing them under different temperatures and mechanical conditions and by approximating the shape of actual cells. Also, we reproduced the cell shape using a cylindrical model this time. However, since the real cell has a complicated shape, it is necessary to use random numbers and precise modeling of the cytoskeleton structure for verification. Further elucidation of the molecular mechanism of cytoskeletal control by the mechanical stimulation received by cells is expected by developing techniques that can measure the force and shape of the cytoskeleton.

All the cells that make up our body receive mechanical stimuli and maintain homeostasis through appropriate haptic responses. Its failure is expected to be the cause of many diseases. By advancing the elucidation of the process by which cells receive mechanical stimuli and transform them into chemical signals, many discoveries have been made about problems that could not be explained only by chemical signals. It is expected that it will be useful in investigating mechanical cell behaviors.

## Conflicts of Interest

The author declares that the present study has no conflict of interest.

## Notes

### Competing Interest Statement

The authors have declared no competing interest.

